# Effects and Mechanisms of Aerobic Exercise on Myocardial AGEs/RAGE-p38 MAPK-NF-*κ*B Pathway in SHR Rats

**DOI:** 10.64898/2026.03.07.710255

**Authors:** Tingting Nong, Shanshan Liu, Xiaogui Pan

## Abstract

**Objective:** To investigate the effects of aerobic swimming exercise on blood pressure, cardiac function, and the myocardial AGEs/RAGE-p38 MAPK-NF-*κ*B signaling pathway in spontaneously hypertensive rats (SHR).

**Methods:** Twenty-four male Wistar-Kyoto (WKY) rats were randomly assigned to three control subgroups (C-0, C-4, C-8; n=8 each). Fifty-six SHRs were allocated into seven subgroups (S-0, S-4, S-8, SE-4, LE-4, SE-8, and LE-8; n=8each) to receive different swimming intervention protocols. Systolic blood pressure (SBP) and diastolic blood pressure (DBP) were measured via the tail-cuff method. Cardiac parameters, including ejection fraction (EF), fractional shortening (FS), left ventricular end-diastolic volume (LVEDV), left ventricular end-systolic volume (LVESV), and stroke volume (SV), were assessed using echocardiography. Myocardial morphological alterations were observed through hematoxylin-eosin (HE) staining, and myocardial hydroxyproline content was quantified using the alkaline hydrolysis method. Furthermore, the myocardial expressions of advanced glycation end products (AGEs), receptor for AGEs (RAGE), phosphorylated p38 mitogen-activated protein kinase (p-p38 MAPK), and nuclear factor-kappa B (NF-*κ*) were detected by immunohistochemistry (IHC).

**Results:** Aerobic exercise significantly reduced blood pressure in SHRs, partially improved cardiac function and myocardial architecture, and decreased hydroxyproline content (except in the SE-4 group). Furthermore, the exercise intervention downregulated the expressions of AGEs, RAGE (except in the SE-4 group), p-p38 MAPK, and NF-*κ*, with efficacy varying according to exercise duration and intervention cycles.

**Conclusion:** Aerobic exercise alleviates the progression of hypertension and mitigates the risk of cardiac dysfunction in SHRs by inhibiting myocardial glycation. This cardioprotective effect may be mediated by the suppression of p38 MAPK activation and the subsequent reduction of NF-*κ* nuclear translocation.

## Introduction

Hypertension, a pervasive and insidious pathological condition, remains one of the most significant cardiovascular threats to global public health. Notably, its global prevalence among adults reached a critical 33% in 2023 [1]. This alarming statistic underscores the exigency of developing effective strategies to mitigate this widespread health crisis.

The pathogenesis of hypertensive complications is closely linked to advanced glycation end products (AGEs). Beyond direct interactions with cell wall proteins, AGEs bind to their specific receptor (RAGE), thereby triggering a cascade of deleterious biological responses. This signaling axis induces the activation of key mediators, including p38 MAPK, NF-*κ*, endothelin-1 (ET-1), intercellular adhesion molecule-1 (ICAM-1), and vascular cell adhesion molecule-1 (VCAM-1) [2]. Persistent activation of these pathways facilitates further elevation of blood pressure and a progressive decline in cardiac function, thereby exacerbating cardiovascular damage [3]. Consequently, attenuating AGE-mediated detrimental effects represents a pivotal therapeutic strategy for preventing hypertension and mitigating associated cardiac remodeling.

Accumulating evidence has established that regular exercise is a potent non-pharmacological intervention for lowering blood pressure and enhancing myocardial performance [4] [5]. However, limited scientific attention has been directed toward the impact of exercise on myocardial AGEs and RAGE expression, particularly concerning the complex molecular mechanisms governing hypertensive progression.

In the present study, we aimed to delineate the effects of distinct exercise regimens on the AGEs/RAGE-p38 MAPK-NF-*κ* axis in SHRs. This research seeks to illuminate the regulatory role of physical activity in these molecular pathways and identify novel therapeutic and preventive avenues for hypertension-related cardiac disorders. Distinct from previous literature, this work systematically investigates the dose-response relationship regarding exercise duration (30 min vs. 60 min) and intervention cycles (4 weeks vs. 8 weeks) on the targeted signaling pathway.

## Materials and methods

### Experimental Animals

Twenty-four male Wistar-Kyoto (WKY) rats and fifty-six male SHRs (age: 8 weeks; body weight: 180–200 g) were purchased from Beijing Vital River Laboratory Animal Technology Co., Ltd. [License No. SCXK (Beijing) 2012-0001]. Animals were housed in a temperature-controlled environment (22 ± 2°C) with a relative humidity of 45% ± 10% and a 12-hour light/dark cycle. Rats were provided with a standard rodent diet and water ad libitum. Following a one-week acclimation period, WKY rats were randomly assigned to three control groups: 0-week control (C-0), 4-week control (C-4), and 8-week control (C-8) (n=8 each). SHRs were randomly allocated into seven experimental subgroups (n=8 each): 0-week (S-0), 4-week (S-4), and 8-week (S-8) sedentary groups; and four exercise intervention groups: 4-week 30-min exercise (SE-4), 4-week 60-min exercise (LE-4), 8-week 30-min exercise (SE-8), and 8-week 60-min exercise (LE-8). The detailed grouping strategy is summarized in Table1.

**Table 1.** Grouping of experimental animals.

| Time point | Control group (n=24) | Sedentary group (n=24) | 30-min exercise group (n=16) | 60-min exercise group (n=16) |
| --- | --- | --- | --- | --- |
| 0 week | C-0 (n=8) | S-0 (n=8) | — | — |
| 4 weeks | C-4 (n=8) | S-4 (n=8) | SE-4 (n=8) | LE-4 (n=8) |
| 8 weeks | C-8 (n=8) | S-8 (n=8) | SE-8 (n=8) | LE-8 (n=8) |

Note: C-0/C-4/C-8: WKY rats in 0/4/8-week control group; S-0/S-4/S-8: SHR rats in 0/4/8-week sedentary group; SE-4/SE-8: SHR rats in 4/8-week 30-min exercise group; LE-4/LE-8: SHR rats in 4/8-week 60-min exercise group; “—” indicates no corresponding group.

### Exercise Intervention Protocol

The WKY control and SHR sedentary groups remained in standard housing without exercise intervention. The SHR exercise groups underwent a swimming training program in a fiber-reinforced polymer pool (150×60×70 cm) with a water depth of 60 cm and a temperature of 32–34°C. After a 1-week adaptive swimming period, a progressive training protocol was initiated: daily swimming duration was incrementally increased by 5–10 minutes per session, starting from 5–10 minutes on the first day and reaching the target duration (30 or 60 min/day) by the sixth day. This target intensity was maintained until the conclusion of the 4-week or 8-week training period. Rats were closely monitored during exercise; those that ceased swimming were gently encouraged with a glass rod. Following each session, rats were immediately dried with sterile towels to prevent hypothermia.

### Blood Pressure and Cardiac Function Assessment

SBP and DBP were measured via the tail-cuff method using the CODA™ 8 multi-channel non-invasive system (Kent Scientific, USA). Cardiac function was evaluated using the Vevo 770™ high-resolution small animal imaging system (VisualSonics, Canada). Rats were anesthetized with a single intraperitoneal injection of 2% sodium pentobarbital (40-60 mg/kg) and were secured on a heating pad (37°C). The precordial region was shaved and an acoustic coupling gel was applied. An RMV-710B scan head was used to obtain 2D left ventricular (LV) long-axis and M-mode images. Functional parameters including EF, FS, LVM, LVEDV, LVESV, and SV were calculated.

### Sample Collection and Biochemical Analysis

#### Tissue Collecting

Myocardial samples were collected 24 hours after the final exercise session. Rats were euthanized via an overdose of sodium pentobarbital (100–150 mg/kg, i.p.). Following the confirmation of deep anesthesia, myocardial tissues were thoroughly perfused with ice-cold normal saline to eliminate residual blood. Harvested samples were either fixed in 4% paraformaldehyde for histological analysis or snap-frozen and stored at -80°C for biochemical assays.

#### Histological and Molecular Detection

Myocardial hydroxyproline content was quantified using the alkaline hydrolysis method. For morphological assessment, tissues were paraffin-embedded, sectioned at 5 µm, and subjected to H&E staining. The protein expressions of AGEs, RAGE, p-p38 MAPK, and NF-*κ* were detected using commercial IHC kits (Elabscience Biotechnology Co., Ltd., China). Six non-overlapping fields per section were analyzed using a Leica image analysis system; results were quantified as the mean optical density (IOD/Area).

#### Histological and Immunohistochemical (IHC) Evaluation

For morphological assessment, myocardial tissues were paraffin-embedded, sectioned at a thickness of 5 µm, and subjected to hematoxylin-eosin (H&E) staining. The sections were examined under a light microscope to evaluate cardiomyocyte architecture and exercise-induced structural remodeling.

To elucidate the underlying molecular mechanisms, the expressions of AGEs, RAGE, p-p38 MAPK, and NF-*κ* were detected using commercial IHC kits (Elabscience Biotechnology Co., Ltd., Wuhan, China). For each section, six non-overlapping representative fields were randomly selected and digitized using a Leica image analysis system to ensure objectivity. The protein expression levels were quantified as the integrated optical density (IOD) normalized to the area (IOD/Area), providing a standardized parameter for intergroup comparisons.

## Statistical Analysis

Statistical analyzes were performed using SPSS Statistics 26.0. Data are presented as mean ± SD. Intergroup differences were analyzed using one-way ANOVA followed by Tukey’s post-hoc test. A P-value <0.05 was considered statistically significant.

## Results

### Effects of Aerobic Exercise on Blood Pressure

As illustrated in Fig1 (adapted from Pan et al. [5]), no significant differences in SBP or DBP were observed among the WKY control subgroups (C-0, C-4, and C-8). In contrast, both SBP and DBP were significantly elevated in the SHR sedentary groups compared with age-matched WKY rats (P<0.05 or P<0.01), with hypertensive progression becoming more pronounced as the rats aged. Compared with their age-matched sedentary counterparts (S-4 and S-8), the exercise intervention groups (SE and LE) exhibited significant reductions in both SBP and DBP. Specifically, in the 4-week intervention cohort, the LE-4 group showed a significantly lower DBP than the SE-4 group (P<0.05), although no such difference was observed for SBP. However, in the 8-week intervention cohort, no significant differences in blood pressure were detected between the SE-8 and LE-8 groups. Furthermore, there were no statistically significant variations in blood pressure levels when comparing the different exercise durations and cycles (SE-4, SE-8, LE-4, and LE-8) against each other.

**Fig 1.**
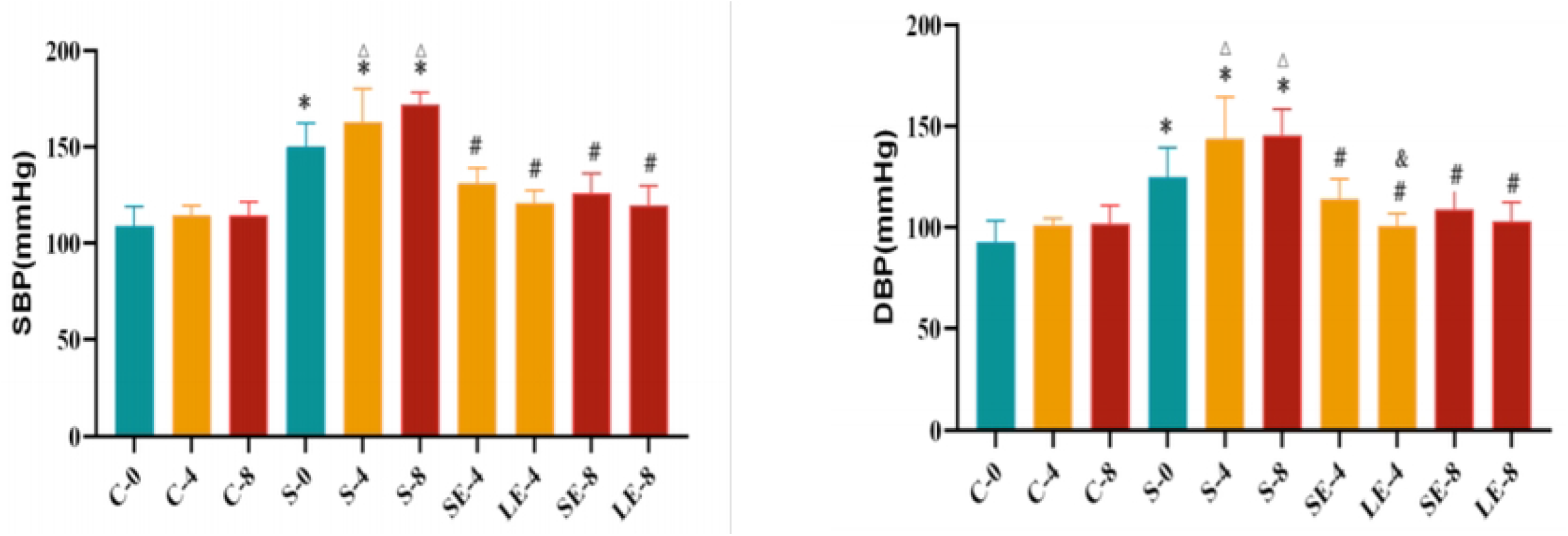
**B**lood pressure. A: Systolic blood pressure (SBP, mmHg) in each group. B: Diastolic blood pressure (DBP, mmHg) in each group. Note: compared with the age-matched control group, *p<0.05; compared with S-0 group, Δ p<0.05; compared with the age-matched sedentary group, #p<0.05; compared with 30min exercise group, &p<0.05.

### Impact of Exercise on Cardiac Function

As summarized in Table 2, the cardiac functional and structural parameters exhibited significant variations across the groups. Compared with the C-0 group, the S-4 group demonstrated a significant decline in EF and FS, while the S-8 group showed a marked increase in LVM.

**Table 2.** Cardiac function parameters in each experimental group.

| Groups | EF (%) | FS (%) | LVM/mg | LVEDV (μL) | LVESV (μL) | SV (μL) |
| --- | --- | --- | --- | --- | --- | --- |
| C-0 | 69.61±3.40 | 41.13±2.11 | 688.11±44.31 | 290.19±22.17 | 88.23±12.19 | 201.96±17.24 |
| C-4 | 67.46±4.66 | 39.21±2.51 | 749.23±70.61 | 276.13±16.37 | 87.48±6.60 | 188.66±19.96 |
| C-8 | 67.46±3.61 | 38.71±2.95 | 770.48±58.24 | 273.89±15.45 | 88.73±6.56 | 185.16±19.39 |
| S-0 | 66.71±4.20 | 40.05±5.56 | 669.51±81.93 | 273.09±29.49 | 91.24±16.74 | 181.85±19.56 |
| S-4 | 64.04±5.26Δ | 36.18±3.93Δ | 750.56±68.03 \$ | 297.12±46.94 | 108.30±30.36 * \$ | 188.82±21.00 |
| S-8 | 68.19±5.42 | 39.50±4.43 | 843.21±103.75ΔΔ * \$\$ | 297.35±24.72 | 94.21±15.56 | 203.14±27.78 |
| SE-4 | 68.90±2.33 # | 44.91±2.91 # # | 748.26±47.15 | 302.08±27.23 | 93.93±10.61 | 208.15±20.59\$ |
| LE-4 | 68.23±3.13 | 44.16±3.01 # # | 785.13±34.02 | 286.52±19.92 | 91.09±11.02 # | 195.43±15.73 |
| SE-8 | 65.72±5.03 | 40.40±7.14 & | 826.96±50.97 | 306.18±39.62\$ | 104.78±20.39 | 201.40±31.19 |
| LE-8 | 69.39±3.14 | 40.36±2.51 | 831.58±34.13 | 297.78±28.85 | 91.62±16.46 | 206.16±16.92\$ |

Note: Compared with S-0 group: \$ P ≤ 0.05; Compared with age-matched control group: \* P ≤ 0.05; Compared with age-matched sedentary group: # P ≤ 0.05, ## P ≤ 0.01; Compared with 4-week exercise group: & P ≤ 0.01.

Intragroup analysis within the SHRs revealed that, relative to the S-0 baseline, the S-4 group exhibited significantly elevated LVM and LVESV, and the S-8 group maintained a heightened LVM. In contrast, aerobic exercise showed compensatory and protective effects: SV was significantly increased in the SE-4 and LE-8 groups, and LVEDV was increased in the SE-8 group compared to the S-0 group.

When compared with age-matched WKY rats, the S-4 group displayed a significantly higher LVESV, and the S-8 groupshowed a significantly increased LVM. Notably, aerobic exercise partially reversed these hypertensive alterations. Compared with the S-4 sedentary group, both EF and FS were significantly improved in the SE-4 group, and FS was enhanced in the LE-4 group. Furthermore, LVESV was significantly reduced in the exercise groups compared to the S-4 group, indicating attenuated ventricular remodeling.

Longitudinal comparison revealed that FS was significantly lower in the SE-8 group than in the SE-4 group. Further statistical analysis indicated no significant interaction effect between exercise duration (30 min vs. 60 min) and the intervention period (4 weeks vs. 8 weeks) on the measured cardiac parameters.

### Myocardial Histological Morphology (H&E Staining)

Histopathological evaluation via H&E staining revealed distinct morphological differences among the groups (Fig2). In the WKY control groups (C-0, C-4, and C-8), the myocardial tissue exhibited a highly organized architectural arrangement, with cardiomyocytes displaying uniform size, clear boundaries, and centrally located nuclei.

**Fig 2.**
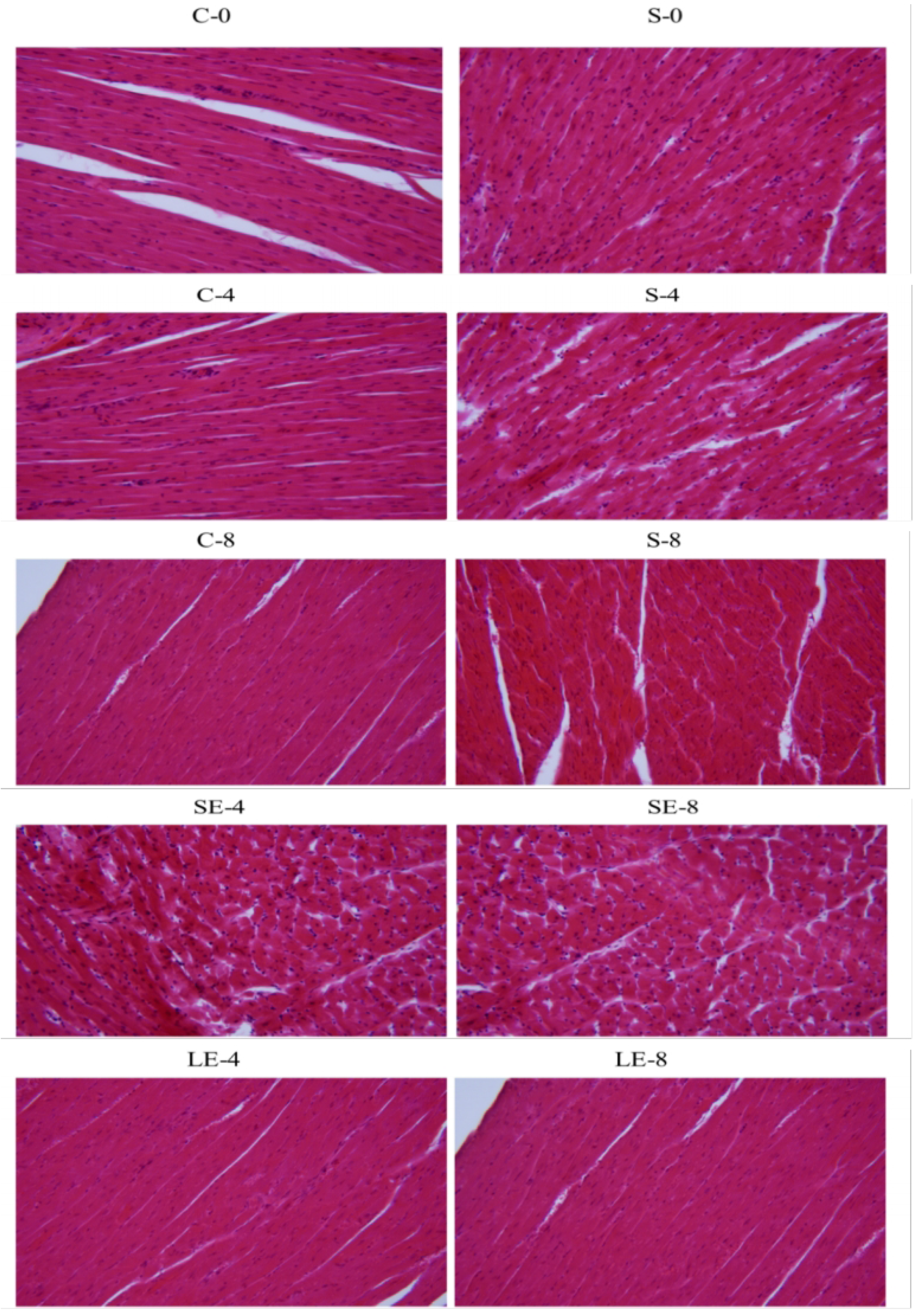
Myocardial hematoxylin-eosin staining. Representative images of myocardial hematoxylin-eosin staining in each group: C-0, C-4, C-8, S-0, S-4, S-8, SE-4, SE-8, LE-4, and LE-8.

In contrast, the SHR sedentary groups (S-4 and S-8) showed prominent pathological remodeling, characterized by marked cardiomyocyte hypertrophy, thickening of myocardial fibers, and interstitial disarray. Furthermore, partial myofilament fragmentation and cytoplasmic vacuolization were observed, indicating progressive hypertensive myocardial injury.

Following the aerobic exercise intervention, the myocardial architecture in the SHR exercise groups (SE and LE) appeared relatively more ordered and regular compared to the sedentary groups. The exercise groups exhibited more distinct cellular boundaries and a visible attenuation of cardiomyocyte hypertrophy. These results suggest that regular swimming exercise effectively preserves myocardial structural integrity and mitigates the hypertrophic response induced by chronic hypertension.

### Myocardial Hydroxyproline Content

The concentration of myocardial hydroxyproline, a key biochemical marker of collagen deposition and fibrosis, is presented in Fig3. No statistically significant differences in hydroxyproline levels were observed among the WKY control subgroups (C-0, C-4, and C-8) or among the SHR sedentary subgroups (S-0, S-4, and S-8).

**Fig 3.**
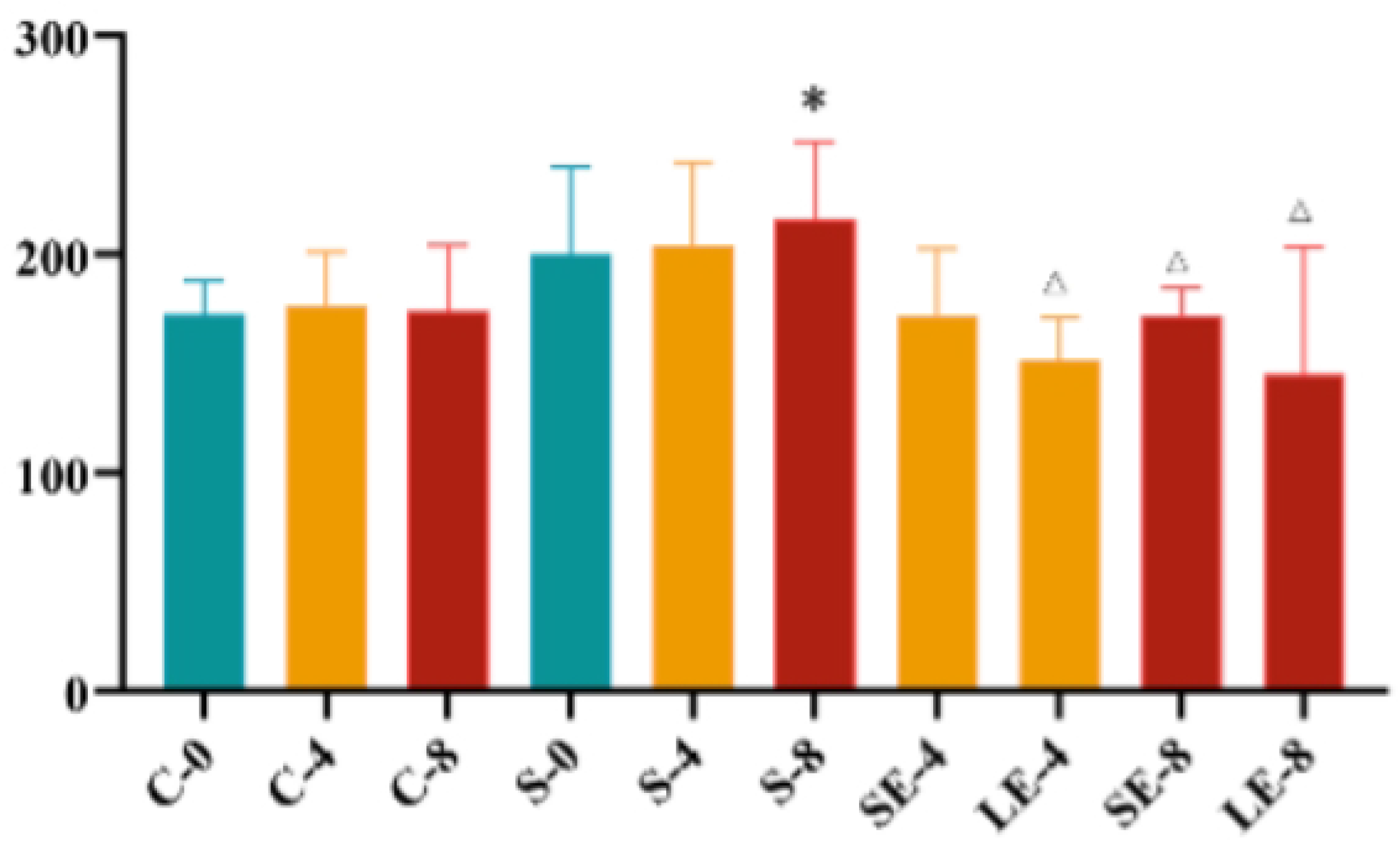
Myocardial hydroxyproline content. Myocardial hydroxyproline content in each experimental group (C-0, C-4, C-8, S-0, S-4, S-8, SE-4, SE-8, LE-4, LE-8). Note: Compared with the control group, *p < 0.05; compared with the age-matched sedentary group, Δp <0.05.

However, when compared with age-matched WKY rats, the S-8 group exhibited a significant elevation in myocardial hydroxyproline content (P<0.05), indicating progressive collagen accumulation associated with chronic hypertension.

Notably, aerobic exercise effectively attenuated this fibrotic process. Compared to their respective age-matched sedentary SHR counterparts, hydroxyproline levels were significantly reduced in the LE-4, SE-8, and LE-8 groups (P<0.05). These findings suggest that both increasing exercise duration (60 min) and extending the intervention period (8 weeks) play a critical role in inhibiting hypertensive myocardial fibrosis.

### Myocardial Expression of AGEs

The protein expression and localization of AGEs in myocardial tissues were evaluated via immunohistochemical (IHC) staining (Fig4 and Fig5). AGE-positive staining was primarily localized within the myocardium cytoplasm, characterized by distinct brown granular deposits (indicated by arrows).

**Fig 4.**
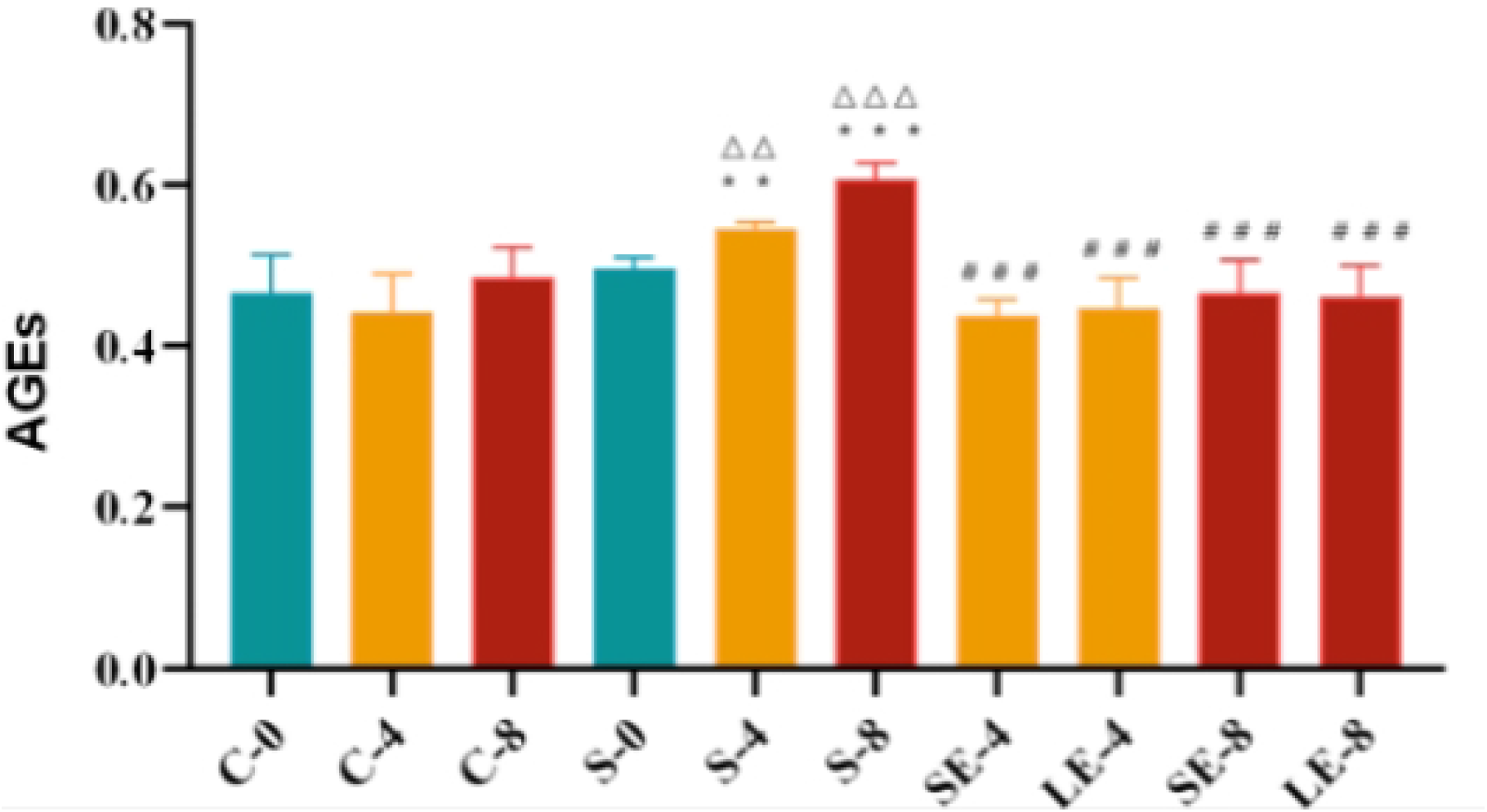
Positive expression of AGEs (IOD/Area) in each group. Positive expression of advanced glycation end products (AGEs, IOD/Area) in each experimental group (C-0, C-4, C-8, S-0, S-4, S-8, SE-4, SE-8, LE-4, LE-8). Note: Compared with the age-matched control group, **p < 0.01; compared with S-0 group, ΔΔp <0.01, ΔΔΔp <0.001; compared with the age-matched sedentary group, ##p< 0.01.

**Fig 5.**
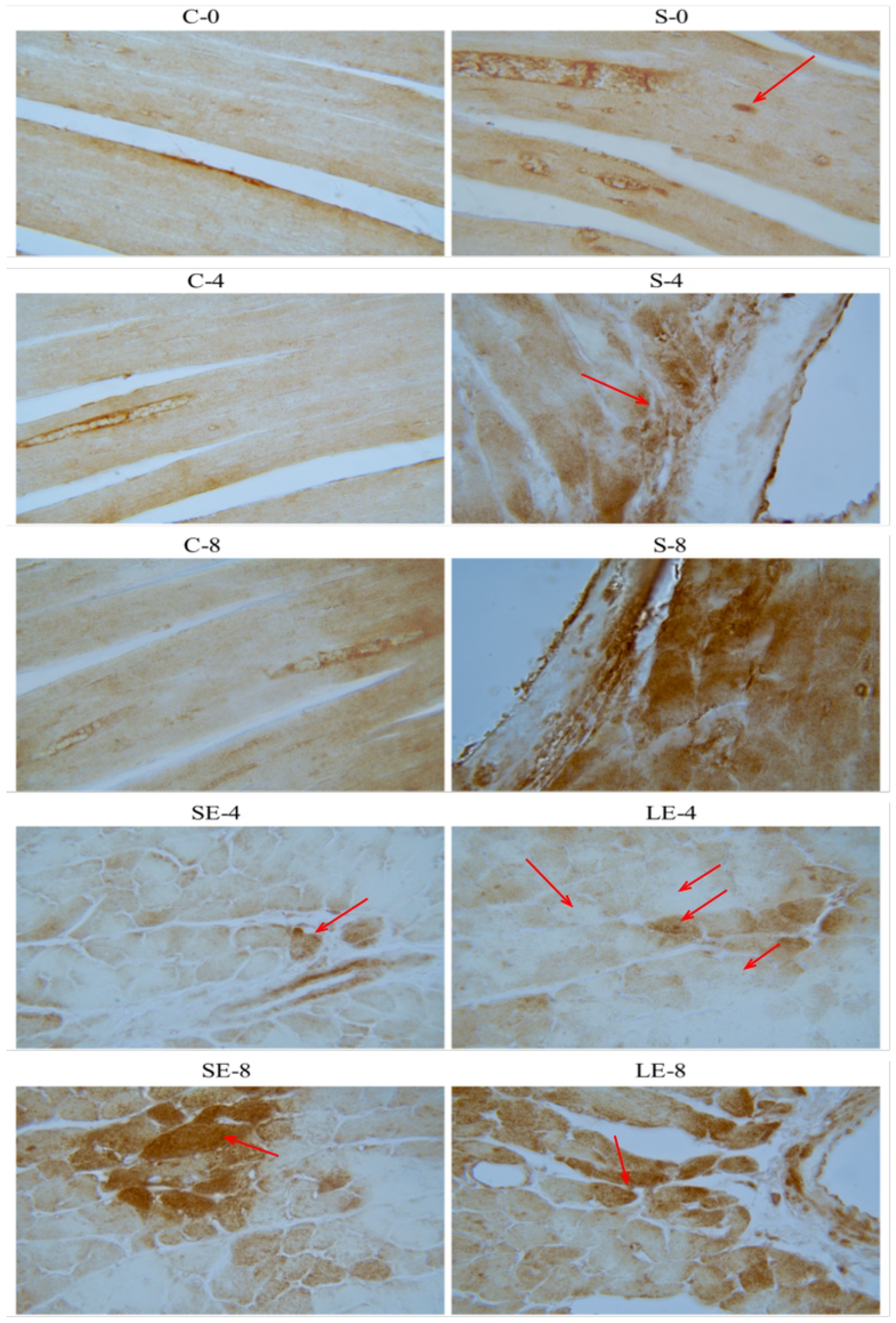
Immunohistochemical staining of AGEs. Representative images of immunohistochemical staining for advanced glycation end products (AGEs) in each group (C-0, C-4, C-8, S-0, S-4, S-8, SE-4, SE-8, LE-4, LE-8). Magnification: 10×100.

Compared with age-matched WKY rats, the SHR sedentary groups (S-4 and S-8) exhibited significantly higher AGE-positive expression than the respective C-4 and C-8 groups (P¡0.05). Within the SHR cohort, AGE expression showed a progressive increase over time: levels in the S-4 group were significantly elevated relative to the S-0 baseline, while the S-8 group demonstrated a marked escalation compared to the S-4 group (P¡0.01), indicating cumulative glycation stress during hypertensive progression.

In contrast, aerobic exercise effectively suppressed the accumulation of myocardial AGEs. Compared with the S-4 group, AGE-positive expression was significantly downregulated in both the SE-4 and LE-4 exercise groups. Similarly, both the SE-8 and LE-8 groups showed a significant reduction in AGE expression compared to the S-8 group (P¡0.05). Notably, no significant differences were detected between the 30-min and 60-min exercise durations, nor between the 4-week and 8-week intervention cycles within the exercise groups, suggesting that even moderate exercise duration is sufficient to yield significant anti-glycation benefits.

### Myocardial RAGE Expression Levels

As illustrated in Fig6 and Fig7, RAGE was predominantly localized on the membranes of cardiomyocytes and vascular endothelial cells. Compared with age-matched WKY rats, the SHR sedentary groups (S-4 and S-8) exhibited significantly higher RAGE-positive expression than the C-4 and C-8 groups, respectively (P<0.05).

**Fig 6.**
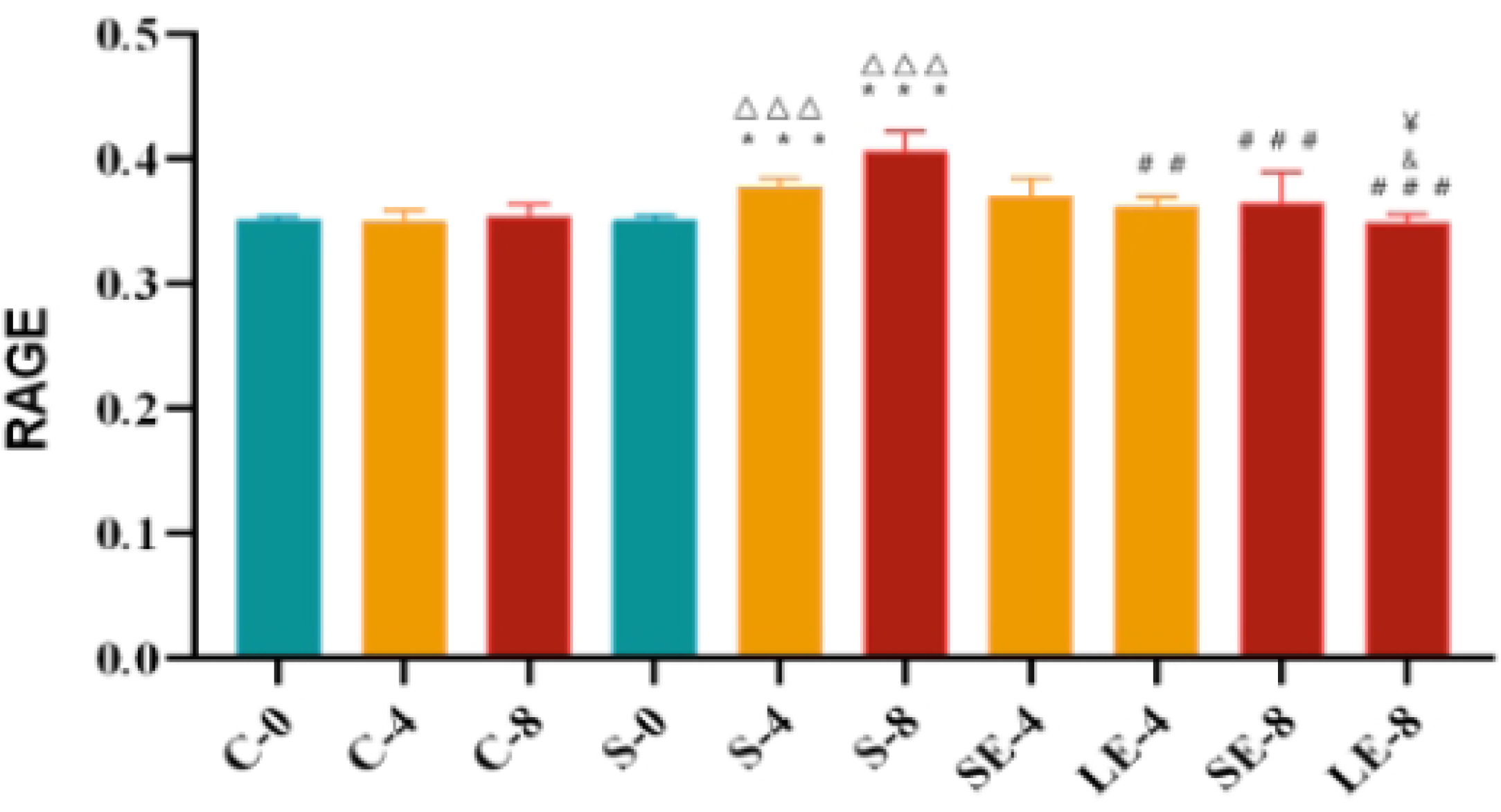
Positive expression of RAGE (IOD/Area) in each group. Positive expression of receptor for advanced glycation end products (RAGE, IOD/Area) in each experimental group (C-0, C-4, C-8, S-0, S-4, S-8, SE-4, SE-8, LE-4, LE-8). Note: Compared with the control, **p<0.001; compared with the S-0 group, ΔΔΔp<0.01; compared with the age-matched group, ##p<0.01; compared with the exercise group at week 4, &p<0.05; compared with the 30-minute exercise group, ¥p < 0.05.

**Fig 7.**
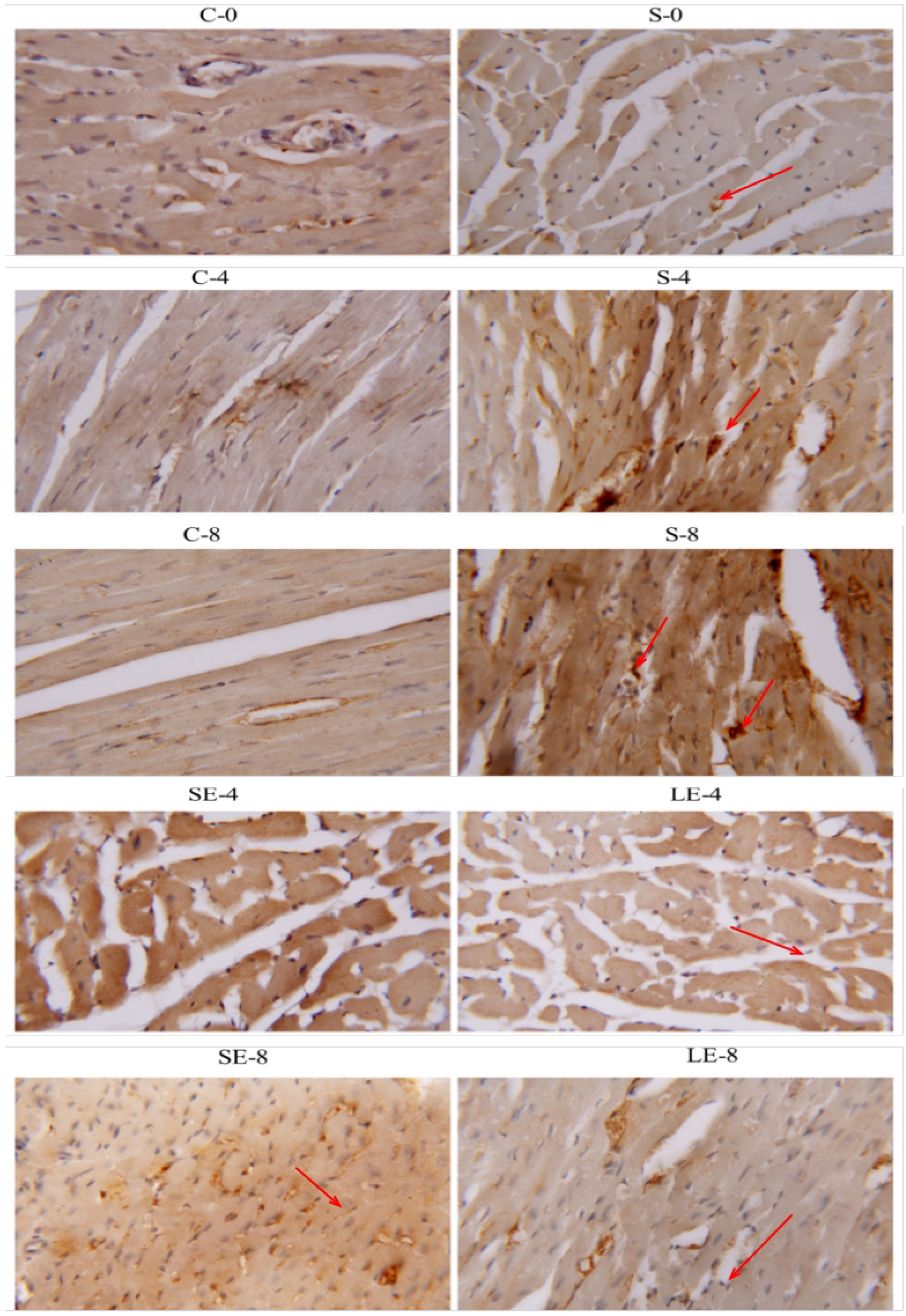
Immunohistochemical staining of RAGE. Representative images of immunohistochemical staining for receptor for advanced glycation end products (RAGE) in each group (C-0, C-4, C-8, S-0, S-4, S-8, SE-4, SE-8, LE-4, LE-8). Magnification: 10×100.

Within the SHR groups, RAGE expression demonstrated a time-dependent escalation: a highly significant increase was observed in the S-4 group compared with the S-0 baseline (P<0.01), with a further, more pronounced elevation in the S-8 group relative to the S-4 group (P<0.01). These data indicate that RAGE expression is profoundly upregulated during the progression of hypertensive myocardial injury.

In contrast, aerobic exercise significantly downregulated RAGE expression. In the 4-week cohort, the LE-4 groupshowed a significant reduction in RAGE-positive expression compared to the S-4 group (P<0.05). In the 8-week cohort, both the SE-8 and LE-8 groups exhibited a highly significant decrease in RAGE expression compared to the S-8 group(P<0.01).

Comparative analysis of different exercise regimens revealed no significant differences in RAGE expression between the SE-4 and LE-4 groups, nor between the SE-8 and LE-8 groups. However, it is noteworthy that RAGE expression in the LE-8 group was lower than that in both the SE-8 and LE-4 groups, suggesting that prolonged, higher-duration exercise might offer incremental benefits in suppressing RAGE-mediated signaling.

### Myocardial p-p38 MAPK Expression Levels

As illustrated in Fig8 and Fig9, the protein expression of phosphorylated p38 MAPK (p-p38 MAPK) was utilized to evaluate the activation status of the p38 signaling pathway. Compared with age-matched WKY rats, the positive expression of p-p38 MAPK was significantly higher in both the S-4 and S-8 sedentary groups than in their respective C-4 and C-8 counterparts (P<0.05).

**Fig 8.**
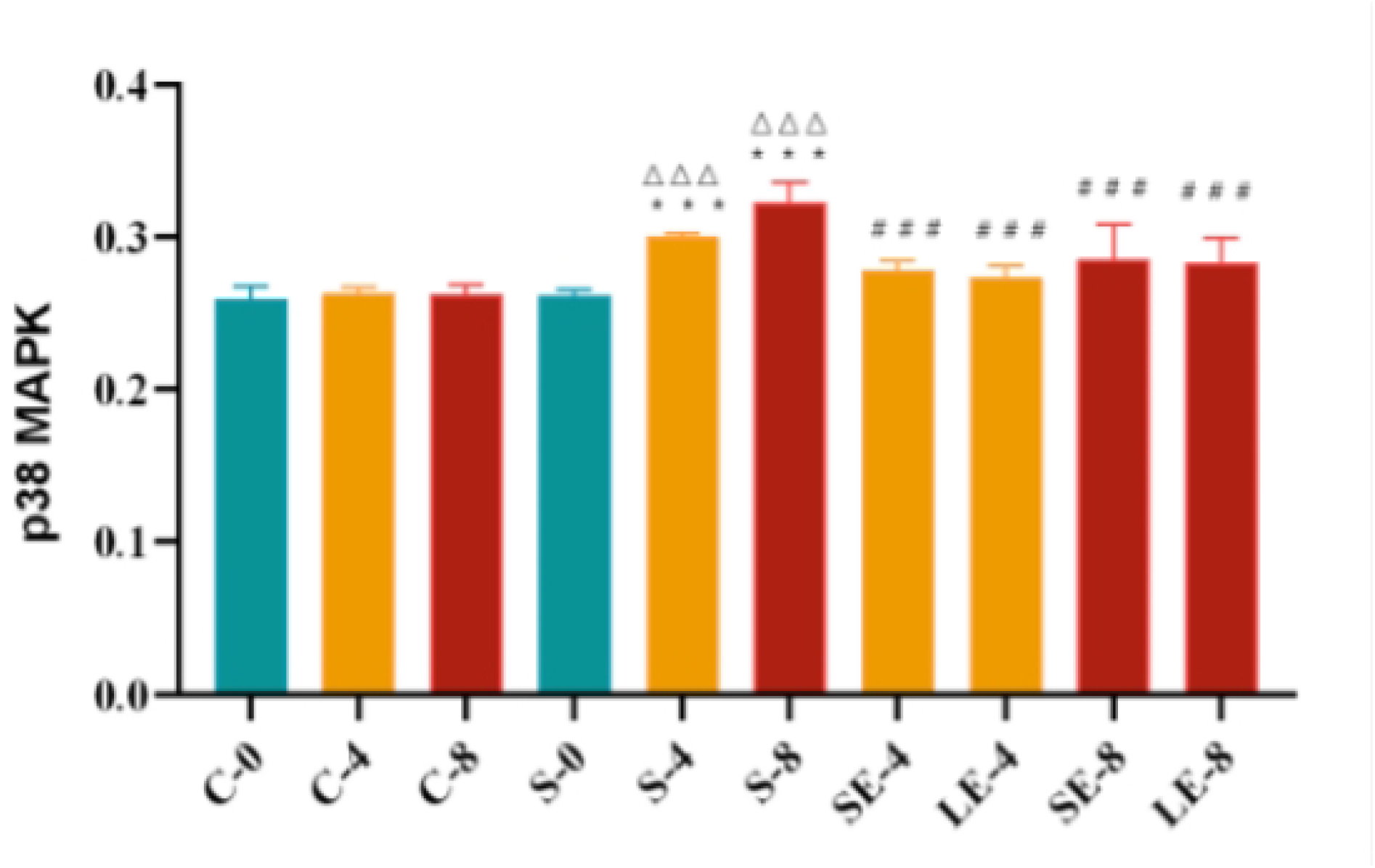
Positive expression of phospho-p38 MAPK (IOD/Area) Positive expression of phosphorylated p38 mitogen-activated protein kinase (phospho-p38 MAPK, IOD/Area) in each experimental group (C-0, C-4, C-8, S-0, S-4, S-8, SE-4, SE-8, LE-4, LE-8). Note: Compared with the age-matched control group, **p<0.01; compared with S-0 group, ΔΔΔp<0.001; compared with the age-matched control group, ###p <0.001.

**Fig 9.**
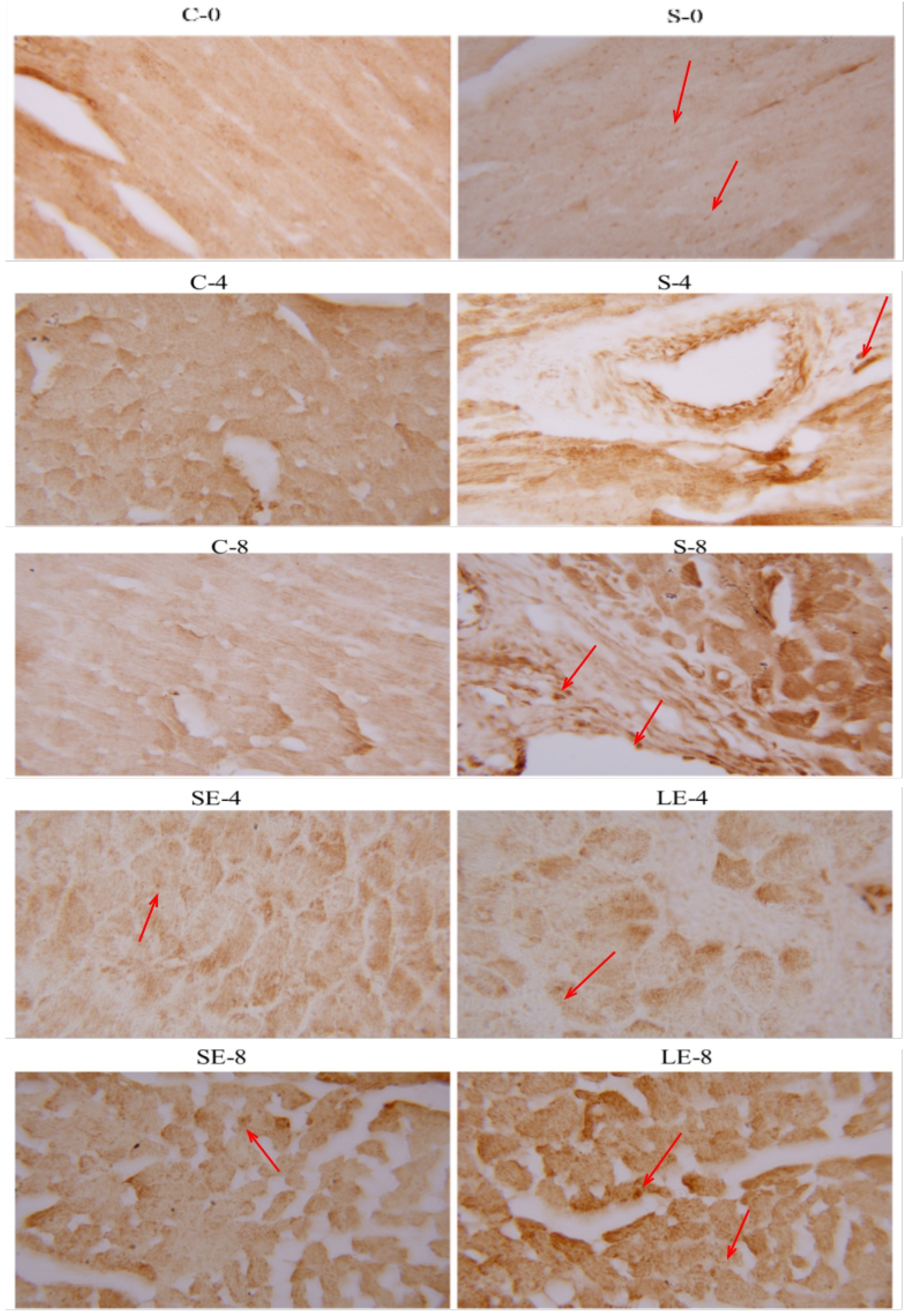
Immunohistochemical expression of phospho-p38 MAPK. Representative images of immunohistochemical staining for phosphorylated p38 mitogen-activated protein kinase (phospho-p38 MAPK) in each group (C-0, C-4, C-8, S-0, S-4, S-8, SE-4, SE-8, LE-4, LE-8). Magnification: 10×100.

Within the SHR cohort, a significant increase in p-p38 MAPK levels was observed in both the S-4 and S-8 groups relative to the S-0 baseline, suggesting that the activation of the p38 MAPK pathway is closely associated with the progression of hypertension.

In contrast, aerobic exercise significantly suppressed p38 MAPK activation. Compared with their respective age-matched sedentary counterparts (S-4 and S-8), p-p38 MAPK expression levels were significantly decreased in all exercise intervention groups (SE-4, LE-4, SE-8, and LE-8; P<0.05). Furthermore, no significant differences in the expression of p-p38 MAPK were detected between the SE-4 and LE-4 groups, nor between the SE-8 and LE-8 groups, indicating that both 30-minute and 60-minute exercise durations were equally effective in attenuating p38 MAPK phosphorylation in this model.

### Myocardial NF-*κ* Expression Levels

The protein expression of NF-*κ*, a downstream transcription factor in the AGEs/RAGE-p38 MAPK pathway, was examined to assess the extent of myocardial inflammatory signaling (Fig10 and Fig11).

**Fig 10.**
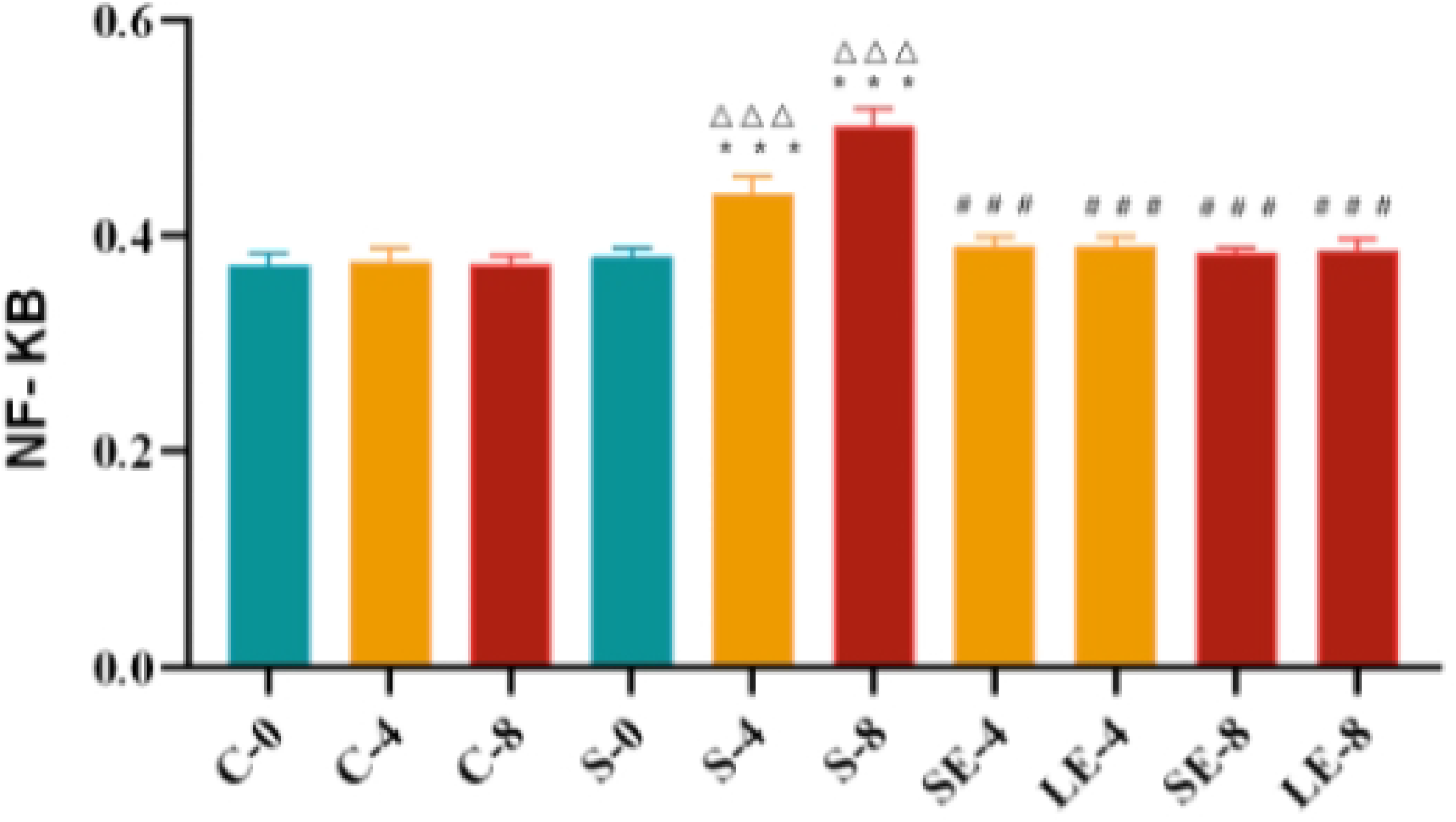
Positive expression of NF-*κ* (IOD/Area) Positive expression of nuclear factor kappa B (NF-*κ*, IOD/Area) in each experimental group (C-0, C-4, C-8, S-0, S-4, S-8, SE-4, SE-8, LE-4, LE-8). Note: Compared with the age-matched control group, **p< 0.01; compared with S-0 group, ΔΔΔp< 0.001; compared with the age-matched control group, ###p < 0.001.

**Fig 11.**
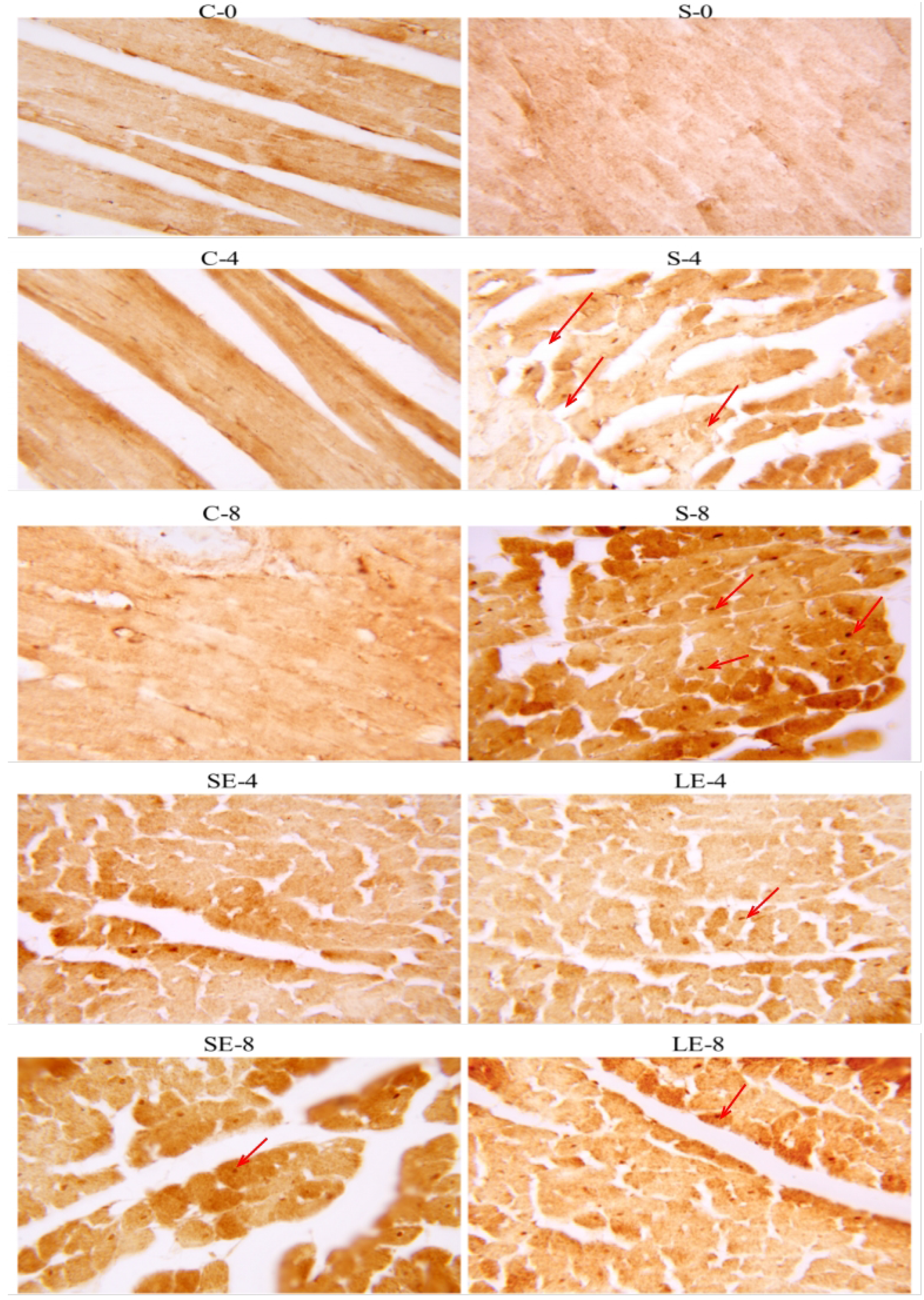
Immunohistochemical staining of NF-*κ*. Representative images of immunohistochemical staining for nuclear factor kappa B (NF-*κ*) in each group (C-0, C-4, C-8, S-0, S-4, S-8, SE-4, SE-8, LE-4, LE-8). Magnification: 10×100.

Compared with age-matched WKY rats, NF-*κ*-positive expression in the S-4 and S-8 groups was highly significantly elevated relative to the C-4 and C-8 groups, respectively (P<0.01). Furthermore, compared with the S-0 baseline, the expression of NF-*κ* showed an extreme significant increase in both the S-4 and S-8 sedentary groups (P<0.01), highlighting the progressive activation of pro-inflammatory transcription in the hypertensive heart.

In contrast, aerobic exercise intervention effectively mitigated NF-*κ* activation. Compared with their respective age-matched sedentary counterparts (S-4 and S-8), the positive expression of NF-*κ* was significantly reduced in all exercise groups (SE-4, LE-4, SE-8, and LE-8; P<0.05). Similar to the findings for p-p38 MAPK, no significant differences in NF-*κ* levels were observed between the SE-4 and LE-4 groups or between the SE-8 and LE-8 groups. These results demonstrate that swimming exercise, regardless of the daily duration (30 vs. 60 min), consistently inhibits the activation of NF-*κ* signaling in SHRs.

## Discussion

The stable blood pressure observed in WKY rats across different time points aligns with previous literature, confirming their validity as normotensive controls. In contrast, SBP and DBP in SHRs escalated significantly with age, exceeding the hypertensive threshold (140/90 mmHg). These findings characterize the model as representative of mild-to-moderate hypertension. According to the WHO Global Report on Hypertension, physical activity is a universally beneficial intervention for hypertensive populations [1]. Consistent with prior studies [5], our data demonstrate that moderate-intensity aerobic swimming effectively attenuates or even reverses the progression of hypertension in SHRs. Notably, we found that both lower-intensity and varied durations of swimming exerted significant antihypertensive effects, reinforcing the potency of exercise as a non-pharmacological strategy.

Pathological remodeling, including myocardial fibrosis and cardiomyocyte hypertrophy, is a hallmark of hypertensive heart disease. While the S-4 group showed a significant decline in EF and FS, these parameters remained relatively stable in the S-8 group, suggesting that left ventricular contractility may not deteriorate linearly with blood pressure elevation. This observation supports the hypothesis that the transition from left ventricular hypertrophy (LVH) to overt heart failure is a protracted process [6]. Our results show that 4 weeks of swimming training improved EF and FS, particularly in the SE-4 group, indicating an early enhancement of myocardial contractility. Further progression of hypertension may eventually lead to significant cardiac hypertrophy [7]. Interestingly, the lack of significant differences in LVM across exercise groups—consistent with Yuan et al. [8]—suggests that while exercise improves function, its impact on established structural hypertrophy may require longer intervention or higher intensity.

Hydroxyproline serves as a reliable biomarker for myocardial interstitial fibrosis. In this study, a significant increase in hydroxyproline content was only observed in the S-8 group, suggesting that myocardial fibrosis in SHRs manifests at later pathological stages. Following 12 weeks of swimming training, myocardial hydroxyproline levels were previously reported to be markedly reduced in diabetic models [9]. Similarly, our study found that swimming protocols of 30 min/8 weeks, 60 min/4 weeks, and 60 min/8 weeks significantly reduced hydroxyproline levels, potentially reversing existing fibrosis. The limited efficacy of the 30 min/4 weeks protocol likely points to a minimum threshold of exercise volume required to stimulate collagen degradation or inhibit its deposition. A central finding of this study is the modulation of the AGEs/RAGE axis. Myocardial AGEs accumulation in SHRs was positively correlated with blood pressure. The cross-linking of AGEs with extracellular matrix proteins like collagen and elastin impairs arterial elasticity and reduces susceptibility to hydrolytic enzymes, thereby increasing arterial stiffness [10]. Furthermore, AGEs can impair vascular function by inhibiting nitric oxide synthase (NOS) activity [11]. Our results demonstrate that swimming exercise significantly reduces myocardial AGEs content. Although the dose-response relationship between 30 and 60 minutes was minimal for AGEs levels, the overall reduction indicates a systemic shift in glycation stress.

The binding of AGEs to RAGE triggers the generation of reactive oxygen species (ROS) and subsequent activation of the p38 MAPK and NF-*κ* pathways. This cascade drives the expression of inflammatory and profibrotic factors, such as VCAM-1, ICAM-1, and PAI-1, promoting cardiovascular damage [12] [13]. We observed that swimming exercise markedly downregulated RAGE expression, potentially through two mechanisms: (1) direct reduction of the AGEs ligand, and (2) potential induction of sRAGE (soluble RAGE), which acts as a decoy receptor to block AGE-RAGE signaling [14] [15]. Notably, the 60 min/8 weeks regimen was the most effective in suppressing RAGE, highlighting the benefits of prolonged intervention.

As a stress-activated protein kinase, p38 MAPK phosphorylation is a critical mediator of cardiac hypertrophy and fibrosis [16]. AGEs can bind to RAGE, leading to p38 MAPK phosphorylation and the activation of various inflammatory pathways, thereby triggering an inflammatory response [17]. Our study confirmed that p-p38 MAPK levels were elevated in SHRs and further reduced by aerobic exercise. Inhibition of p38 MAPK is increasingly recognized as a central mechanism of cardioprotection [18] [19]. Similarly, the downregulation of NF-*κ*, a downstream effector of p38 MAPK, further supports the anti-inflammatory role of exercise. Previous studies have shown that increased oxidative stress and impaired autophagy significantly elevate p-NF-*κ* and P62 levels in cardiomyocytes, whereas aerobic exercise for more than two months effectively reduces these markers [20]. Furthermore, in an aging rat model of diet-induced atherosclerosis, 8 weeks of swimming training was shown to significantly reduce myocardial NF-*κ* expression [21].By reducing NF-*κ* activation, swimming training likely mitigates the “cytokine storm” and oxidative stress-induced autophagy impairment often seen in the hypertensive myocardium.

In conclusion, our study demonstrates that swimming exercise alleviates hypertensive myocardial injury in SHRs. This protective effect is mediated, at least in part, by the inhibition of the AGEs/RAGE-p38 MAPK-NF-*κ* signaling pathway, providing a robust molecular basis for the use of aerobic exercise in hypertension management.

## Conclusion

In summary, aerobic exercise alleviates the progression of hypertension and mitigates the risk of cardiac dysfunction in SHRs by suppressing the myocardial AGEs/RAGE signaling axis. This cardioprotective mechanism is closely associated with the inhibition of p38 MAPK phosphorylation and the subsequent reduction of NF-*κ* nuclear translocation. Notably, while swimming exercise demonstrated potent therapeutic efficacy, extending the daily exercise duration (from 30 to 60 min) or the intervention period (from 4 to 8 weeks) did not result in further significant improvements in blood pressure or the measured molecular markers. These findings suggest that moderate aerobic exercise protocols are sufficient to yield substantial anti-glycation and anti-inflammatory benefits, providing experimental evidence for exercise as an effective non-pharmacological strategy for managing hypertensive myocardial injury.

